# *In vivo* effect of recombinant FSH and LH administered to meagre (*Argyrosomus regius*) at the initial stages of sex differentiation

**DOI:** 10.1101/2023.12.30.573690

**Authors:** Álvaro González Cid, Ignacio Giménez, Neil Duncan

## Abstract

Recombinant gonadotropins, follicle stimulating (rFSH) and luteinizing hormone (rLH), offer the potential to induce gametogenesis in prepubertal fish. This study aimed to determine the *in vivo* effect of *Argyrosomus regius* rFSH (arrFSH) and *Argyrosomus regius* rLH (arrLH) administered to prepubertal meagre juveniles at the initial stages of sex differentiation. Meagre single-chain recombinant gonadotropins, arrFSH and arrLH were produced with the CHO expression system by Rara Avis Biotec, S. L. Juvenile meagre, 9-months old with mean weight of 222 ± 36 g (mean ± SD) were randomly distributed into seven experimental groups (n = 8 per group) that were treated weekly for three weeks with an acute injection of 6, 12 or 18 μg kg^-1^ of arrFSH (groups, 6-arrFSH, 12-arrFSH and 18-arrFSH) or 6, 12 or 18 μg kg^-1^ of arrLH (groups, 6-arrLH, 12-arrLH and 18-arrLH) or saline solution (Control group). Two more groups (n=8) were set up and treated for 6 weeks, with 12 μg kg^-1^ of arrFSH or saline Control. The fish were held in a 10 m^3^ tank with natural photoperiod (Feb. – March) and temperature 16.1 ± 0.4°C. At the start of the experiment (n = 8) and end of the three week experiment all fish were sacrificed and gonads dissected. Gonads were weighed, fixed in Bouin’s solution and processed for histological analysis. Blood was sampled from all fish at the start and end of the experiment (week 3 and 6) for 17β-estradiol (E2) and 11-ketotestosterone (11-KT) analysis. Juvenile meagre at the start of the experiment were in the initial stages of sexual differentiation, indicated by the presence of the ovarian or testes lumen that was surrounded by undifferentiated embryonic germ stem cells and somatic cells. At the end of the experiment, there was no significant difference in gonadosomatic index (GSI) amongst control (initial and saline treated) and the experimental groups. After three weeks of application of arrFSH, arrLH or saline all fish presented a similar gonadal structure as at the start of the experiment. However, the incidence of isolated developing germ cells (principally spermatogonia, spermatocytes, spermatids, but also perinucleolar stage oocytes) generally increased in arrGTH treated meagre. A mean of 44 % of meagre treated with arrFSH or arrLH presented isolate developing germ cells, mainly male cells. Plasma steroid levels of E2 decreased significantly from the start of the experiments to the end. At the end of the experiment there were no differences in plasma E2 amongst Control fish and rGTH treated fish. Plasma 11-KT showed no change from the start of the experiment to week 3. However, a significant increase was observed in the arrFSH group after six weeks of treatment compared to the start of the experiment and the control group on week 6. The application of arrFSH or arrLH to meagre at the initial stages of sex differentiation did not stimulate steroid production until week six and had a limited, but evident effect on the development of isolated germ cells. The rGTHs, arrFSH or arrLH did not stimulate large developmental changes in undifferentiated gonads.

**HIGHLIGHTS:** Exogenous recombinant gonadotropins administered to meagre did not advance sexual differentiation.

Exogenous recombinant gonadotropins administered to meagre did not affect plasma estradiol.

Exogenous recombinant FSH increased the proportion of meagre with isolated male germ cells.

Exogenous recombinant FSH increased plasma 11-ketotestosterone in meagre treated for 6 weeks.

## 1.1 INTRODUCTION

The meagre (*Argyrosomus regius*) is a marine and migratory teleost fish species, which belongs to Sciaenidae family that can be found in the Mediterranean and Black Sea, and along the Eastern coast of the Atlantic Ocean (Kružić et al., 2016; Ramos-Júdez et al., 2019). Meagre have emerged as an important Mediterranean aquaculture species with the third highest finfish production after gilthead seabream (*Sparus aurata*) and European seabass (*Dicentrarchus labrax*). Total Mediterranean production of meagre was 54.917 t in 2022 (APROMAR, 2023). Positive attributes for aquaculture include adaptation to culture systems (Monfort, 2010; Duncan et al., 2013a), good growth reaching 1 kg in the first year (Monfort, 2010; Duncan et al., 2013a; Kružić et al., 2016), good feed conversion ratio (FCR) (0,9-1,2) (Monfort, 2010) (1,8) (Duncan et al., 2013a), relatively easy juvenile production and controlled egg production in captivity (Duncan et al., 2013a; Mylonas et al., 2016).

Female meagre have an asynchronous oocyte development with clear modes for group-synchronous spawners (Duncan et al., 2013a; Gil et al., 2013) and in most cases fail to undergo oocyte maturation, ovulation and spawning in captivity (Duncan et al., 2012, 2013a; Mylonas et al., 2013). Males have an unrestricted, cystic and lobular type testis (Gil et al., 2013) and undergo spermatogenesis, but a proportion of males produce low volumes of sperm in captivity (Fakriadis et al., 2020). Therefore, the synthetic agonist of gonadotropin-releasing hormone (GnRHa) has been used to enhance spermiation (Ramos-Júdez et al., 2019; Fakriadis et al., 2020) and induce oocyte maturation and spawning in females, (Duncan et al., 2012, 2013a; Mylonas et al., 2013; Fernandez-Palacios et al., 2014).

The meagre is a gonochoristic species that reached puberty in captivity at 2+ years in males and 3+ years in females (Schiavone et al., 2012). Size at puberty in a group of 3-year-old fish reared in captivity and successfully used for spawning had mature females > 5.4 kg and males > 4.2 kg (Duncan et al., 2013a) compared to 0.92 ± 0.08 kg in 2 year old males and 1.61 ± 0.09 kg in three year old females in a group of captive meagre that were not used to spawn as breeders (Schiavone et al., 2012). Breeders used in aquaculture are spawned at a large size (> 4 kg) and there is interest to reduce the size and age at puberty for selective breeding programs. Selective breeding programmes enable an increase in production due to a higher growth rate, a reduction of production costs as a result of an improved FCR, a reduction in stress and mortality due to domestication and improved disease resistance (Gjedrem, 2012; Houston et al., 2020). A key factor that determines the time needed to benefit from these genetic advances is the length of the generation interval (Houston et al., 2020). Species that have large generation times such as groupers, tunas or sturgeons take many years to attain puberty and, therefore, the benefits of breeding programs over a fixed time period are greatly reduced. Meagre breeders used in breeding programs tend to be larger than 3 – 4 kg or 3 - 4 years old (Duncan et al., 2013a), which will slow down progress of using breeding programs to increase production characteristics of the cultured stock.

Reproduction and puberty are controlled by the brain-pituitary-gonad (BPG) axis, which in turn is controlled by environmental factors (Duncan et al., 2013b; Taranger et al., 2010). Within this system, the pituitary gonadotropins (GTHs) and their receptors are the key to conveying the hormonal signals released by the BPG axis (Yaron and Levavi-Sivan, 2011). The GTHs are two heterodimeric glycoproteins, follicle stimulating hormone (FSH) and luteinizing hormone (LH) composed of a common alpha subunit linked non-covalently to a gonadotropin specific specific beta subunit that is also species specific. Follicle stimulating hormone stimulates the early stages of gametogenesis, including spermatogenesis in males and oogenesis in females through the respective synthesis of 17β-estradiol (E2) and 11-ketotestosterone (11-KT) via the membrane-bound receptors FSHra, which belongs to the family of G-protein-coupled receptors (GPCRs) (Lubzens et al., 2010; Schulz et al., 2010; Yaron and Levavi-Sivan, 2011). In males, FSHra is expressed in Sertoli and Leydig somatic cells; and in females in the follicular somatic cells of granulosa and theca as well as in connective tissue. Therefore, FSH control the initiation of puberty and increases in FSH, the associated increases in steroids and the initiation of gametogenesis, where perinuclear oocytes progress to cortical alveolus and initiate vitellogenesis and spermatogonia progress to spermatocytes are clear indications that puberty has been attained and maturation is progressing (Taranger et al., 2010).

However, before puberty can initiate the gonads must first complete the process of sexual differentiation and development of the associated somatic cellular structure. The ovaries should develop perinuclear oocytes associated to follicular somatic cells, Granulosa and Theca, through folliculogenesis and the testis spermatogonia associated to Sertoli and Leydig somatic cells. The meagre began sexual differentiation at 5 months with the differentiated gonads present at 10 and 12 months of age (Schiavone et al., 2012). Sexual differentiation has been shown to be controlled by the expression of genes and enzymes related to steroid production and the production of E2 and 11-KT (Nakamura et al., 1998; Devlin and Nagahama, 2002). Morphological changes with the development of a lumen space (ovarian or testis) and oogonia or spermatogonia have been closely associated with an increase in E2 in fish that differentiate into females and an increase in 11-KT in fish that differentiate as males (Nakamura et al., 1998; Nagahama, 1999). The use of exogenous E2 and androgens applied before and during sex differentiation has long been known to induce development as females or males respectively (Devlin and Nagahama, 2002; Guiguen et al., 2010) and studies inhibiting or knocking out steroid production have also confirmed the roles of steroids (Li, 2019). An increase in E2 was observed to coincide with sexual differentiation in the meagre (Schiavone et al., 2012). However, the upstream control of the steroid production during sexual differentiation are in the early stages of being defined. Follicle stimulating hormone has been accepted as the controlling hormone to initiate progression of gametogenesis and associated steroidogenesis in the secondary growth phase of gametogenesis (Lubzens et al., 2010; Schulz et al., 2010) and consequentially FSH has also been implicated as the hormone that controls steroidogenesis in the differentiating gonads (Devlin and Nagahama, 2002; Lubzens et al., 2010). Increases in FSH and related genes and receptors coinciding with steroid production and sexual differentiation have been described in different fish species (Fan et al., 2003; Moles et al., 2011; Fan et al., 2022) and the application of porcine FSH to orange-spotted grouper (*Epinephelus coioides*) induced sexual differentiation (Huang et al., 2019). However, medaka (*Oryzias latipes*), with an FSH receptor knockout phenotype, appeared to undergo sexual differentiation and develop as males and females, although females had small ovaries and infertile gametes compared to males that developed normally (Murozumi et al., 2014). Additionally, immunoreactivity against FSHb and LHb (beta protein of the gonadotropins) was not detected in the pituitary of the Malabar grouper, (*Epinephelus malabaricus*) during sexual differentiation suggesting that pituitary gonadotropins do not play a major role in sex differentiation (Murata et al., 2012). Consequentially, the role of FSH in sexual differentiation remains unclear (Huang et al., 2019) or is different in different fish species (Murozumi et al., 2014).

The use of exogenously administered pure species specific gonadotropins (GTH) has recently become a possibility with the production of recombinant gonadotropins (rGTH). Recombinant gonadotropins have been produced from the cDNA sequences of the gonadotropin that is cloned in expression vectors (plasmids or viruses) that are subcloned in heterologous prokaryotic or eukaryotic systems (Molés et al., 2020). A wide range of expression systems have been used for rGTH production and the Chinese hamster ovary (CHO) cells have been found to produce rGTH glycosylated proteins with terminal sialic acids that have high biological activity in vivo (Molés et al., 2020). The rGTHs have been successfully used to overcome reproductive dysfunctions in adult cultured fish and rFSH has proved to stimulate spermatogenesis in Senegalese sole (*Solea senegalensis*) (Chauvigné et al., 2017, 2018), European eel (*Anguilla anguilla*) (Peñaranda et al., 2018), flathead grey mullet (*Mugil cephalus*) (Ramos-Júdez et al., 2021, 2022) and meagre (Zupa et al., 2023) and oogenesis in flathead grey mullet (Ramos-Júdez et al., 2021, 2022). The entire process of spermatogenesis from spermatogonia through to motile spermatozoids and the entire process of oogenesis from perinuclear oocytes through to viable ovulated ova was induced using a combination of rFSH and rLH administered to adult flathead grey mullet arrested in the early stages of gametogenesis (Ramos-Júdez et al., 2021, 2022).

The aim of this study was to assess the effects of the rGTHs, rFSH and rLH on prepubertal 9-months old meagre, with the hypothesis that *Argyrosomus regius* recombinant follicular stimulating hormone (arrFSH) and *Argyrosomus regius* recombinant luteinizing hormone (arrLH) would induce the progress of maturation from the point of sexual differentiation.

## 1.2 METHODS

### 1.2.1 Fish and Experimental design

A group of pre-pubertal immature meagre were set up in IRTA. These meagre were obtained from the research centre Estação Piloto de Piscicultura de Olhão (EPPO) / Aquaculture Research Station (Olhão, Portugal), which is part of the Instituto Portugués do Mar e da Atmosfera / Portuguese institute for the Ocean and Atmosphere (IPMA) (Lisboa, Portugal). The meagre hatched in IPMA from a spawn on the 22nd May 2020 and were reared to 200g. The meagre arrived in IRTA research facilities in La Rápita (Tarragona, Spain) on the 26th November 2020 and the transport and acclimation appeared to go well. A total of 89 juvenile meagre fish (captivity reared fish) were used that were almost 9-months old and weighed 222 ± 36 g (mean ± SD) when the experiment initiated on the 15th February 2021 under natural photoperiod and temperature. For identification, fish was implanted with a PIT tag (Trovan, Spain). Fish were fed to satiety six days a week with commercial feed (BroodFeed, Sparos, Portugal) and were kept under natural photoperiod. The mean temperature during the experiment was 16,1 ± 0,4°C based on previous experiments with meagre (Duncan et al., 2013a). Juvenile fish were held in a rectangular 10 m^3^ fiber glass tank (2 m × 5 m × 1 m depth) with a biomass of 15,1 kg connected to a recirculation system (IRTAmar®).

The effect of arrFSH and arrLH on immature fish was examined in an experiment performed over three and six weeks, from 15th of February to 8th of March (3 weeks) or 30th of March (6 weeks). Seven groups of fish were set up, plus an initial group that was sacrificed at the start of the experiment. All groups were made up of 8 fish, except for groups Control Saline and arrFSH12 that had 16 fish or two groups of eight (Table 1.).

**Table 1.** Treatment group names, number of fish per group (n) and brief description of the groups treatment.

| Group | Hormone | Dose | n | Description of treatment |
| --- | --- | --- | --- | --- |
| Control 0 | None | 0 $\mu\text{g/kg}$ | 8 | Fish sacrificed at start |
| Saline | None | 0 $\mu\text{g/kg}$ | 16 | Control: weekly saline injections for 3 and 6 weeks |
| arrFSH-6 | arrFSH | 6 $\mu\text{g/kg}$ | 8 | Weekly injection 6 $\mu\text{g/kg}$ of arrFSH for 3 weeks |
| arrFSH-12 | arrFSH | 12 $\mu\text{g/kg}$ | 16 | Weekly injection 12 $\mu\text{g/kg}$ of arrFSH for 3 and 6 weeks |
| arrFSH-18 | arrFSH | 18 $\mu\text{g/kg}$ | 8 | Weekly injection 18 $\mu\text{g/kg}$ of arrFSH for 3 weeks |
| arrLH-6 | arrLH | 6 $\mu\text{g/kg}$ | 8 | Weekly injection 6 $\mu\text{g/kg}$ of arrLH for 3 weeks |
| arrLH-12 | arrLH | 12 $\mu\text{g/kg}$ | 8 | Weekly injection 12 $\mu\text{g/kg}$ of arrLH for 3 weeks |
| arrLH-18 | arrLH | 18 $\mu\text{g/kg}$ | 8 | Weekly injection 18 $\mu\text{g/kg}$ of arrLH for 3 weeks |

Control saline was injected intramuscularly with saline solution, groups arrFSH were injected intramuscularly with arrFSH at the respective dose, 6, 12 or 18 μg kg^-1^ and groups arrLH were injected with arrLH at the respective dose, 6, 12 or 18 μg kg^-1^. The treatments were administered each week on Monday (Tables 2 and 3). Blood samples were taken before treatment injection on days 0, 18 and 43. At the start of the experiment, eight fish were randomly sacrificed as a control group (Control 0). At the end of the three weeks on day 22 all treated fish were sacrificed, with the exception of 8 fish in each of the groups Control Saline and arrFSH12. These two groups were injected with the respective treatment for three more weeks (total 6 weeks) and on week 6 (groups Saline 6W and arrFSH-12 6W) all fish were blood sampled and three fish from each group were sacrificed. For all sampling procedures, fish were anaesthetised with 70 mg L^−1^ of MS-222. To sacrifice fish before tissue collection, fish were overdosed with anaesthesia pithed to destroy the brain. Fish weight, length, gonad weight and liver weight were registered. The gonadosomatic and hepatosomatic indices (GSI and HIS, respectively) were calculated as total organ weight divided by total fish weight mutiplied by 100. Blood and gonad tissue samples were taken from all sacrificed fish. All gonad samples were processed for histology and all blood samples were analyzed for steroids.

**Table 2.** Timeline for 3 week experiment and sampling applied to all groups. Number of weeks, date on Monday when treatments were applied, initial letter of each day of the week, treatment day marked with X and green when, saline, arrFSH or arrLH was applied at the doses indicated in Table 1, Blood, the day blood was extracted for steroid analysis is marked with an X and red, Sacrifice, day when fish were sacrificed is marked with an X and yellow.

| Week | 1 |  |  |  |  |  |  | 2 |  |  |  |  |  |  | 3 |  |  |  |  |  |  | 4 |  |  |  |  |  |  |
| --- | --- | --- | --- | --- | --- | --- | --- | --- | --- | --- | --- | --- | --- | --- | --- | --- | --- | --- | --- | --- | --- | --- | --- | --- | --- | --- | --- | --- |
| Date Monday | 15/2/2021 |  |  |  |  |  |  | 22/2/2021 |  |  |  |  |  |  | 1/3/2021 |  |  |  |  |  |  | 8/3/2021 |  |  |  |  |  |  |
| Day | S | M | T | W | T | F | S | S | M | T | W | T | F | S | S | M | T | W | T | F | S | S | M | T | W | T | F | S |
| Treatment |  | X |  |  |  |  |  |  | X |  |  |  |  |  |  | X |  |  |  |  |  |  |  |  |  |  |  |  |
| Blood |  | X |  |  |  |  |  |  |  |  |  |  |  |  |  |  |  |  |  | X |  |  |  |  |  |  |  |  |
| Sacrifice |  | X |  |  |  |  |  |  |  |  |  |  |  |  |  |  |  |  |  |  |  |  |  | X |  |  |  |  |

**Table 3.**
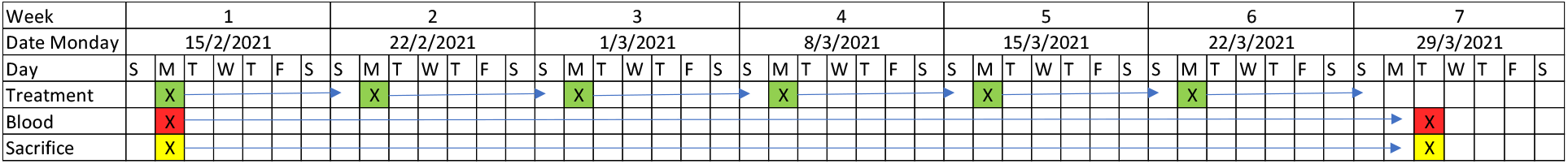
Time line for 6 week experiment and sampling applied to groups Control Saline, and arrFSH12. Number of weeks, date on Monday when treatments were applied, initial letter of each day of the week, treatment - day marked with X and green when, saline of arrFSH was applied at the doses indicated in Table 1, Blood - the day blood was extracted for steroid analysis is marked with an X and red, Sacrifice - day when fish were sacrificed is marked with an X and yellow.

### 1.2.2 Recombinant Gonadotropins

Standard Argyrosomus regius gonadotropin subunit amino acidic sequences (alpha common, beta-FSH and beta-LH) were deduced from RNAseq (Illumina and Nanopore) analysis of mRNA extracted from meagre pituitaries. The mRNA sequences are available in the European Nucleotide Archive (ENA) with project accession number PRJEB57583. The arrFSH sequences were previously reported by Zupa et al., (2023) and the sequences were:

Gonadotropin common alpha subunit

MKRELCLSMVTPATTMGSVKSAGLSLLLLSFLIYVAESYPNIELSNMGCEECTLRKNSVFSRD RPVYQCMGCCFSRAYPTPLKAMKTMTIPKNITSEATCCVAKHSYEIEVAGIRVRNHTDCHCST CYFHKI

*Argyrosomus regius* FSH beta subunit

MQLVVMAAVLAVAGAWQGCGFDCHPTNISIPVESCGNTEFIETTICAGQCYHEDPVYIGHDDWVEQRTCNGDWSYEVKHIKGCPVGVTYPVARNCKCTACNAGSTYCGRFPGDVSSCLSF

*Argyrosomus regius* LH beta subunit

MAIRVSRVMFPLMLTLFLGASSFIWPLAPAVASQLPPCQLINQTVSLEKEGCPKCHPVETTICSGHCITKDPVIKIPFSNVYQHVCTYRDLHYKTFELPDCPPGVDPTVTYPVALSCHCGRCAMDTSDCTFESLQPNFCMNDIPFYY

RARA AVIS BIOTECH S.L. (Valencia, Spain, www.raraavis-bio.com) produced the single-chain *Argyrosomus regius* arrFSH and arrLH. The single-chain sequence used was the beta subunit (FSHβ or LHβ) linked to the 28 amino acids from the carboxyl-terminal of the human chorionic gonadotropin β subunit (hCGβ) linked to the common GTH alpha subunit (FSHα or LHα). The Chinese hamster ovary (CHO) expression system was used to produce the two rGTHs. The CHO culture transfected with the sequence was cultured for 120 h. The cultured rGTHs were purified by ion exchange chromatography and concentrated to 12 μg mL^−1^. Polyclonal (mouse) antibodies for the arrFshβ or arrLHβ subunits and metal affinity purified His-tagged arrFsh or arrLH were used as standards in a semiquantitative Western blot to confirm the concentration.

### 1.2.3 Plasma sex steroid analysis

Blood samples that were collected from caudal vein were placed in Eppendorf tubes and centrifuged at 3000 rpm for 5 min at 4°C; and plasma was aliquoted 3 times (replicas for sex steroid hormones analysis) and stored at -80°C. Plasma steroid levels of 17β-Estradiol (E2) and 11 ketotestosterone (11-KT) were determined by commercial enzyme inmunosorbent assay (EIA, Cayman chemical company, USA). Free steroids were extracted from plasma with methanol. The pellet resulting from evaporation (steroids) was resuspended in ELISA buffer and the steroids E2 and 11-KT were measured in the plasma by ELISA following manufactures instructions (Cayman, Labclinics, Spain).

### 1.2.4 Histological analysis

The effects of the treatments on gonad maturation was evaluated through histological of the gonads and the semi-quantitative analysis of male germ cell proliferation and apoptosis. From each fish, 1-cm thick gonad slices were cut and fixed in Bouin’s solution. Subsequently, gonad samples were dehydrated in ethanol, clarified in xylene, embedded in paraffin wax. Three-μm thick sections were cut and stained with haematoxylin-eosin (H-E). Gonadal tissue structure was observed under a light microscope (Leica DMLB, Houston, USA). Germ cells developmental stage was classified according to the relative size, appearance of structures and morphological changes. The gonadal development of prepubertal meagres was established by evaluating the presence of germ cells in six random optical areas at 20x magnification, although a maximum gonadal development was also established by evaluating the entire gonad due the low presence of sex specific germ cell stages of development. Female germs cells were classified as: chromatin nucleolar stage of primary growth (PGcn) characterized by small oocytes with a large single nucleolus surrounded by a thin layer of citoplasm; perinuclear stage of primary growth (PGps) with a bigger oocyte due to the enlargement of the nucleus and the appearance of multiple nucleoli (West, 1990; Ramos-Júdez et al., 2023). Male germs cells were classified as: type A undifferentiated spermatogonia (StgAund) characterized by being the largest male germ cells and having a large nucleus in addition to one or two nucleoli; type A differentiated spermatogonia (StgAdiff), smaller than StgAund and present in cysts in groups of 2 to 8 germs cells linked by cytoplasmic bridges; type B spermatogonia (StgB) with a smaller size than the previous cells and present in cysts in groups of 16 or more germs cells; spermatocyte (spc) characterized by an increase in cell and nucleus size with respect to StgB due to entry into meiosis (condensing chromosomes), moreover an increase in germs cells per cyst; spermatid (spd) with a significant reduction in cell size and increase of germs cells per cyst; and spermatozoa (spz) with the presence of a flagellum and reduction in cell size (Leal et al., 2009; Schulz et al., 2010; Lacerda et al., 2014).

### 1.2.5 Statistical analysis

Shapiro-Wilk and Levene tests were used to check the normality of data distribution and variance homogeneity, respectively. A one-way analysis of variance (ANOVA) followed by the Holm-Sidak test for pairwise comparisons was used to compare the GSI, HIS, E2, 11-KT and T amongst treatments and sample dates. The Chi-square test was used to compare amongst the treatment groups the ratios of meagre in differentiation sexual with isolated sex specific germ cells. Analyses were performed using SigmaPlot version 12.0 (Systat Software Inc., Richmond, CA, USA). All the results are presented as means ± sd, and the statistical probability significance was established at the P < 0.05 level.

### 1.2.6 Animal experimentation and ethics

The study was conducted in accordance with the European Union, Spanish and Catalan legislation for experimental animal protection (European Directive 2010/63/EU of 22 September on the protection of animals used for scientific purposes; Spanish Royal Decree 53/2013 of February 1st on the protection of animals used for experimentation or other scientific purposes; Boletín Oficial del Estado (BOE), 2013; Catalan Law 5/1995 of June 21th, for protection of animals used for experimentation or other scientific purposes and Catalan Decree 214/1997 of July 30th for the regulation of the use of animals for the experimentation or other scientific purposes). The present study was approved by IRTA’s (Institute of Agrifood Research and Technology) Committee of Ethics and Experimental Animal (CEEA) and the Catalan Government as experimental project 11264 with expedient number FUE-2020-01809522 and ID CJQX0B0PH. The authors complied with the ARRIVE guidelines.

## 1.3 RESULTS

### 1.3.1 Gonadal stage and histology

The group means for GSI ranged from 0.03 ± 0.01 % in group arrFSH-18 to 0.24 ± 0.17 % in group arrFSH-12 6W (Fig. 5.). There were no differences amongst the groups Control 0 and all groups sampled after three weeks of treatment. However, both groups, Saline 6W and aarFSH-12 6W that were treated for six weeks had significantly higher GSI on week 6, compared to Control 0 and all groups sampled at three weeks. The mean group HSI ranged from 1.1 ± 0.2 % in group arrFSH-12 6W to 1.8 ± 0.3 % in group Control 0 (Fig. 6.).

**Figure 5.**
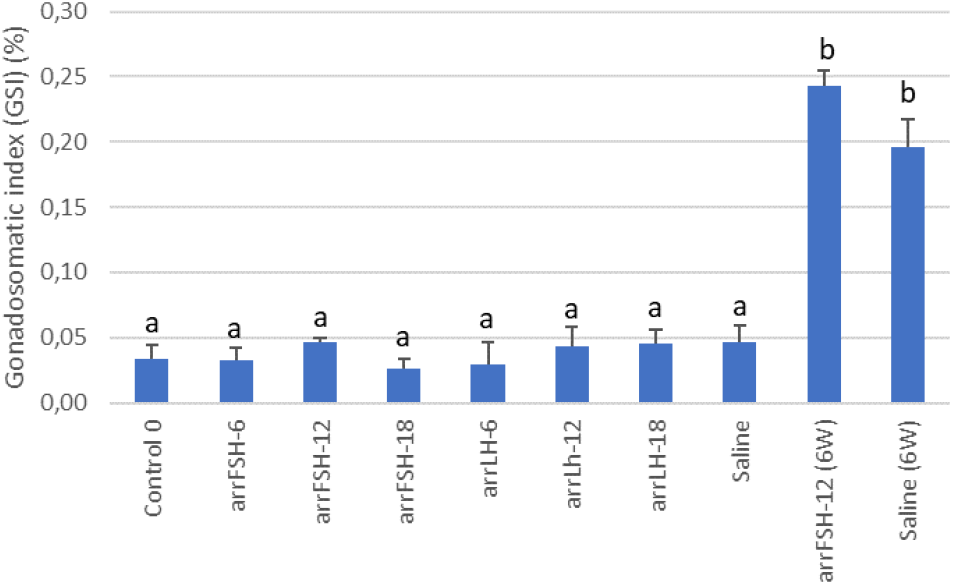
Effect of arrGTH treatments on GSI of meagre during sexual differentiation. Data are the mean ± SD. Different letters denote significant differences (P < 0.05) amongst treatment groups.

**Figure 6.**
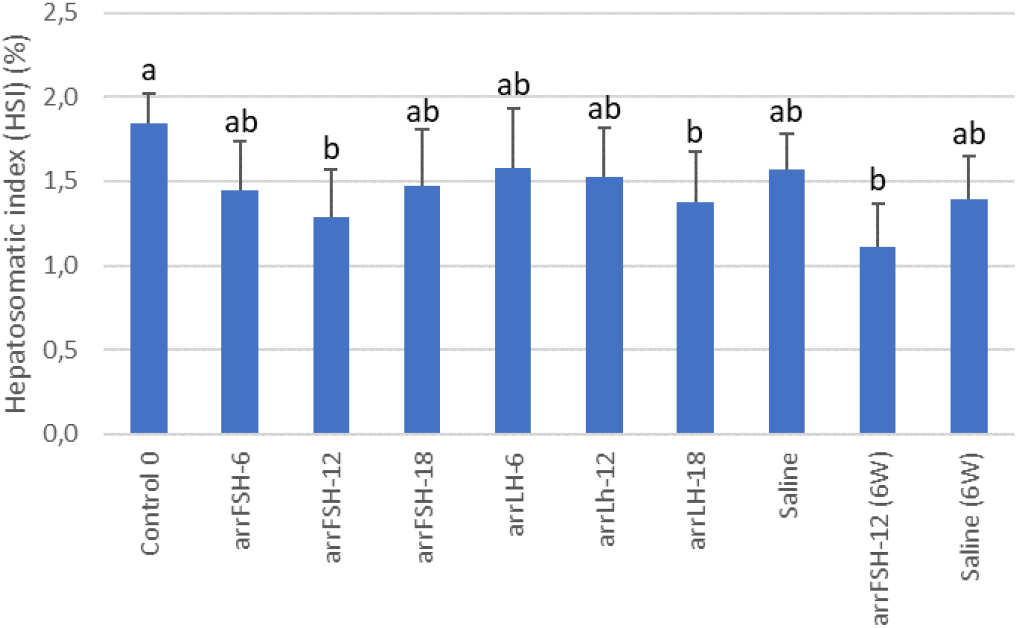
Effect of arrGTH treatments on HSI of immature meagre. Data are the mean ± SD. Different letters denote significant differences (P < 0.05) amongst treatment groups.

The HSI declined significantly from group Control 0 after three weeks in groups arrFSH-12 and arrLH-18 and after 6 weeks in group arrFSH-12. There were no differences amongst the other groups.

The gonads had initiated sexual differentiation and presented the space of the duct or cavity in the centre of the gonad. The duct indicated sexual differentiation had initiated, but gonads had not differentiated (Fig. 7). Moreover, most cells were completely undifferentiated either somatic cells or germ cells that could not be identified as oogonia or spermatogonia. Therefore, all fish sampled were classified as a stage having initiated, but not completed sexual differentiation. The arrGTH treatments increased the presence of highly localised and isolated sexually differentiated germ cells, primary oocytes and type A and B spermatogonia, spermatocytes or spermatids (Fig. 7.).

**Figure 7.**
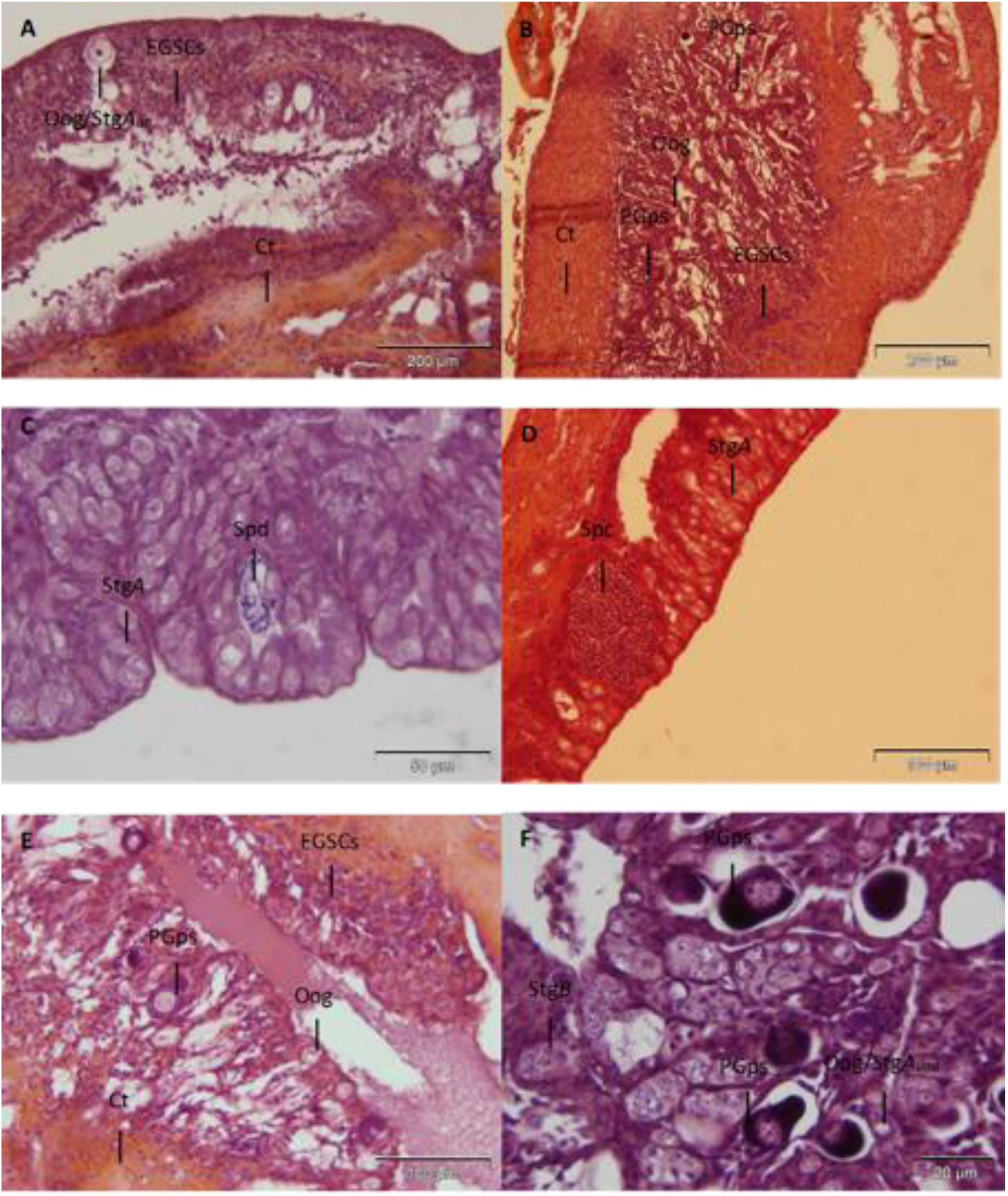
Representative photomicrographs of histological sections of meagre during differentiation sexual, stained with hematoxylin and eosin that were sampled after three weeks of arrGTH treatment. A) sexually undifferentiated fish; B) fish with isolated female germ cells; C) fish isolated male germ cells; D) fish isolated male germ cells; E) fish with isolated female germ cells; F) fish isolated female and male germ cells. EGCS, Embryonic Germ Stem Cells; Oog/Stg, Oogonia or spermatogonia; Oog, oogonia; StgAund, type A undifferentiated spermatogonia; StgB, type B spermatogonia; Spc, spermatocytes; Spd, spermatids; PGps, oocyte at perinucleolar stage of primary growth; Ct, connective tissue.

The arrGTHs significantly increased (P<0.05) the proportion of fish in treated groups with either male or female isolated germ cells compared to control groups (Fig. 8.). Male germs cells (type A and B spermatogonia, spermatocytes or spermatids) were the most common and a proportion of fish treated with arrGTHs were intersex with both male and female germ cells. Groups arrFSH-6 and arrFSH-12 had the highest proportion of fish with gonads that had male or both male and female (intersex) cells, 75 and 62.5 % respectively (Fig. 8.). Other arrGTH treatments varied between 25 and 50% with isolated male or female germ cells. Group arrLH-12 was the only group with gonads that only presented perinuclear oocytes (3 fish or 37.5% of the group). The control groups, Control 0 and control saline had 25% and 0 % respectively of fish with male germ cells. No control fish had the presence of primary oocytes. The fish sampled after 6 weeks presented similar stages of development. One fish from arrFSH-12 6W had female germ cells and one fish from Saline 6W had male germ cells, while other fish had undifferentiated germ cells. Germ cell development was highly localized, most of the gonadal tissue did not have any development and all fish were classified as Stage I (incompletely differentiated).

**Figure 8.**
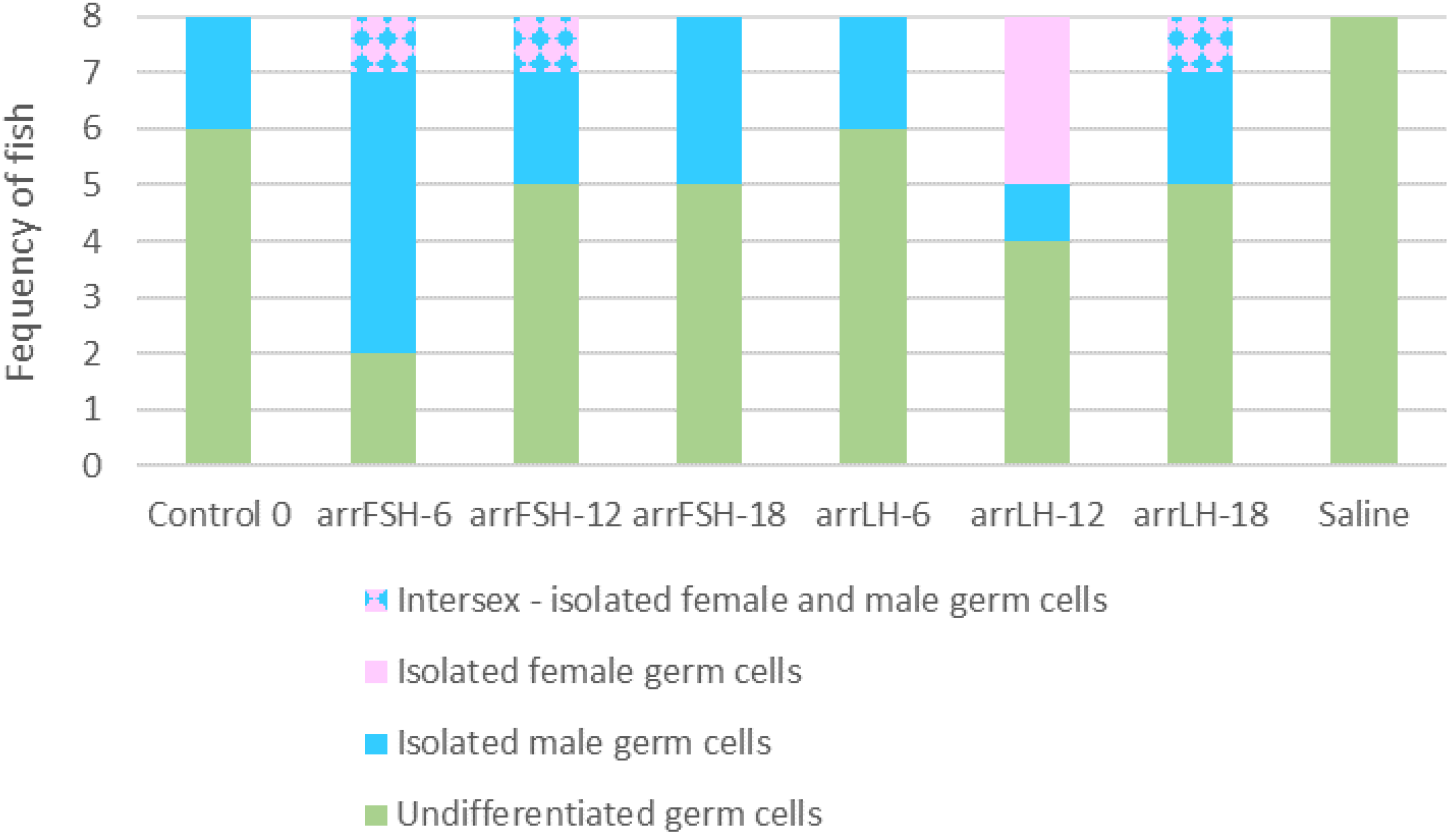
Frequency or number of fish per treatment group that exhibited isolated sexually differentiated germ cells. Undifferentiated germs cells indicates fish in which no differentiated germ cells were observed; Isolated male germ cells, fish with one or more spermatogonia Stg B or higher; Isolated female germ cells, fish with one or more oocytes; Intersex – isolated female and male germ cells, fish that displayed both one or more spermatogonia Stg B or higher and one or more oocytes in the histological cuts.

### 1.3.2 Plasma sex steroids

The plasma steroids were generally not elevated by the arrGTH treatments. Plasma E2 levels decreased significantly (P<0.05) from the start of the experiment, Control 0 (351 ±123 pg ml^-1^) on week 0 to means that ranged from 55 ± 23 pg ml^-1^ in group arrLH-18 to 91 ± 41 pg ml^-1^ in group arrLH-12 (Fig. 9.). After six weeks of treatment the situation was similar and both the means from the groups Saline W6 (138 ± 47 pg ml^-1^) and arrFSH-12 W6 (156 ± 54 pg ml^-1^) were significantly (P<0.05) lower than the Control 0 group (Fig. 10.). No differences were observed in plasma 11-KT plasma levels after 3 weeks of treatment and values ranged from 10 ± 6 pg ml^-1^ in group Saline to 39 ± 28 pg ml^-1^ in group arrFSH-12 (Fig. 11.). After six weeks of treatment a significant (P<0.05) increase in plasma 11-KT was observed in the arrFSH-12 W6 group treated with arrFSH compared to both the Control 0 group (10 ± 3 pg ml^-1^) on week 0 and the Saline control group (8 ± 3 pg ml^-1^) on week 6 (Fig. 12.).

**Figure 9.**
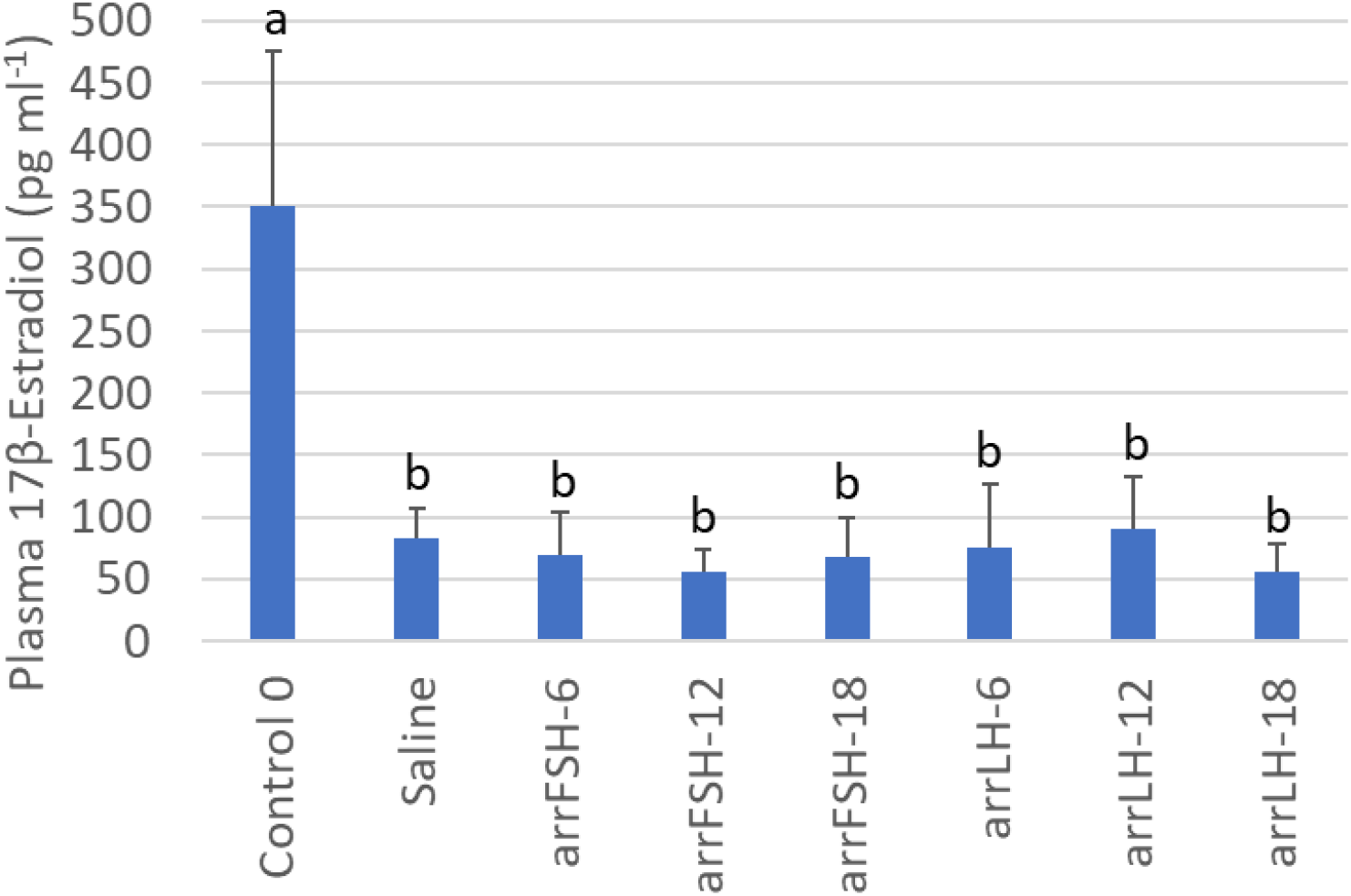
Mean plasma 17β-Estradiol levels of meagre during sexual differentiation in arrGTH treatment groups and Control 0 at the start of the experiment (Week 0) and 4 days after the third weeks injection of the treatment (Week 3) of arrGTH or saline. Different letters indicate a significant (p<0.05) difference between week 0 and 3. The n for Control 0 was 16 and ranged from 6-8 for the different treatment groups.

**Figure 10.**
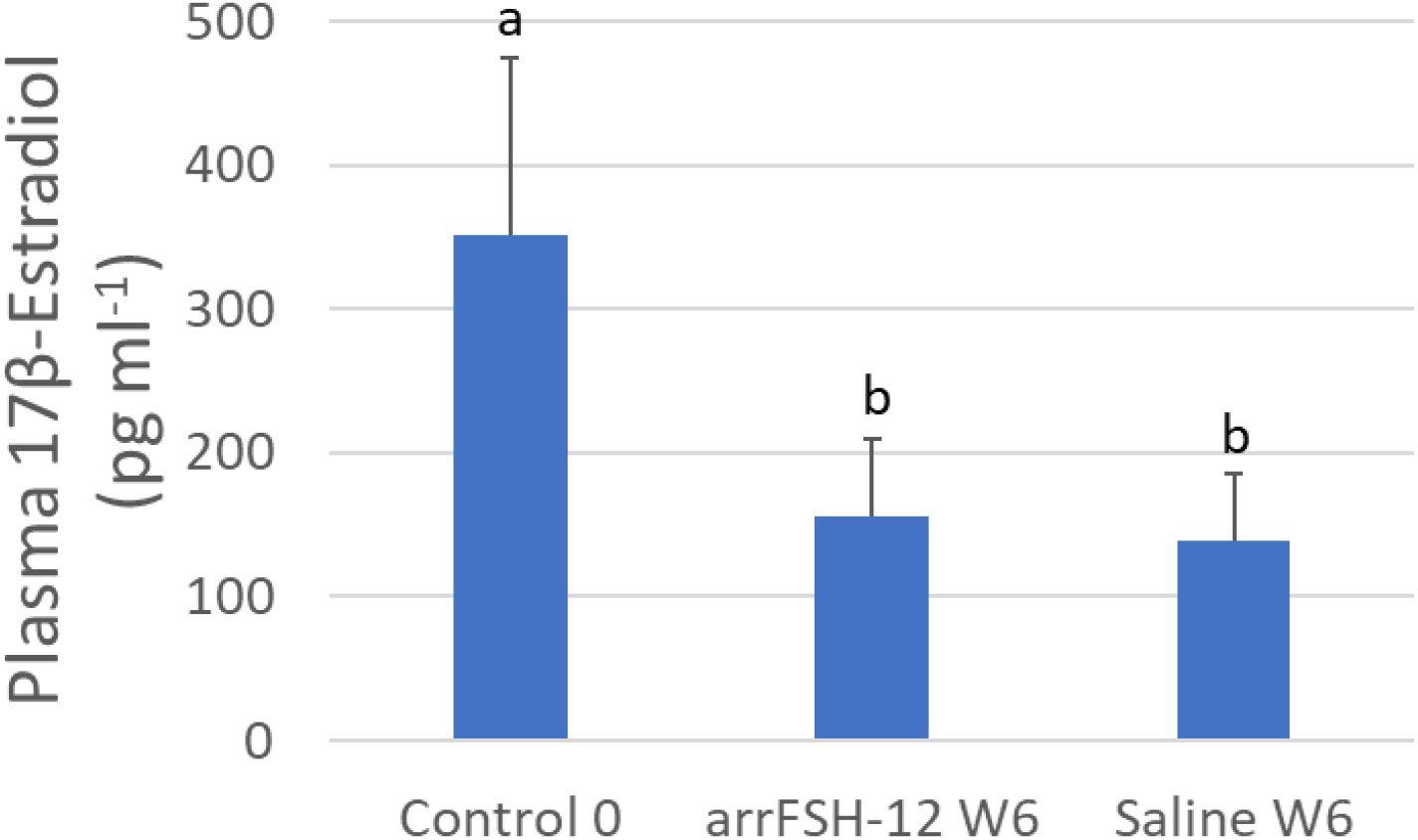
Mean plasma 17β-Estradiol levels of meagre during sexual differentiation at the start of the experiment (Week 0) Control 0 and 7 days after the sixth weeks injection (Week 6) with 12 μg kg^-1^ of arrFSH (arrFSH-12 W6) or Saline (SalineW6 group). Different letters indicate a significant (p<0.05) difference between week 0 and 6. The n for Control 0 was 16 and 8 for groups arrFSH-12 W6 or SalineW6 group.

**Figure 11.**
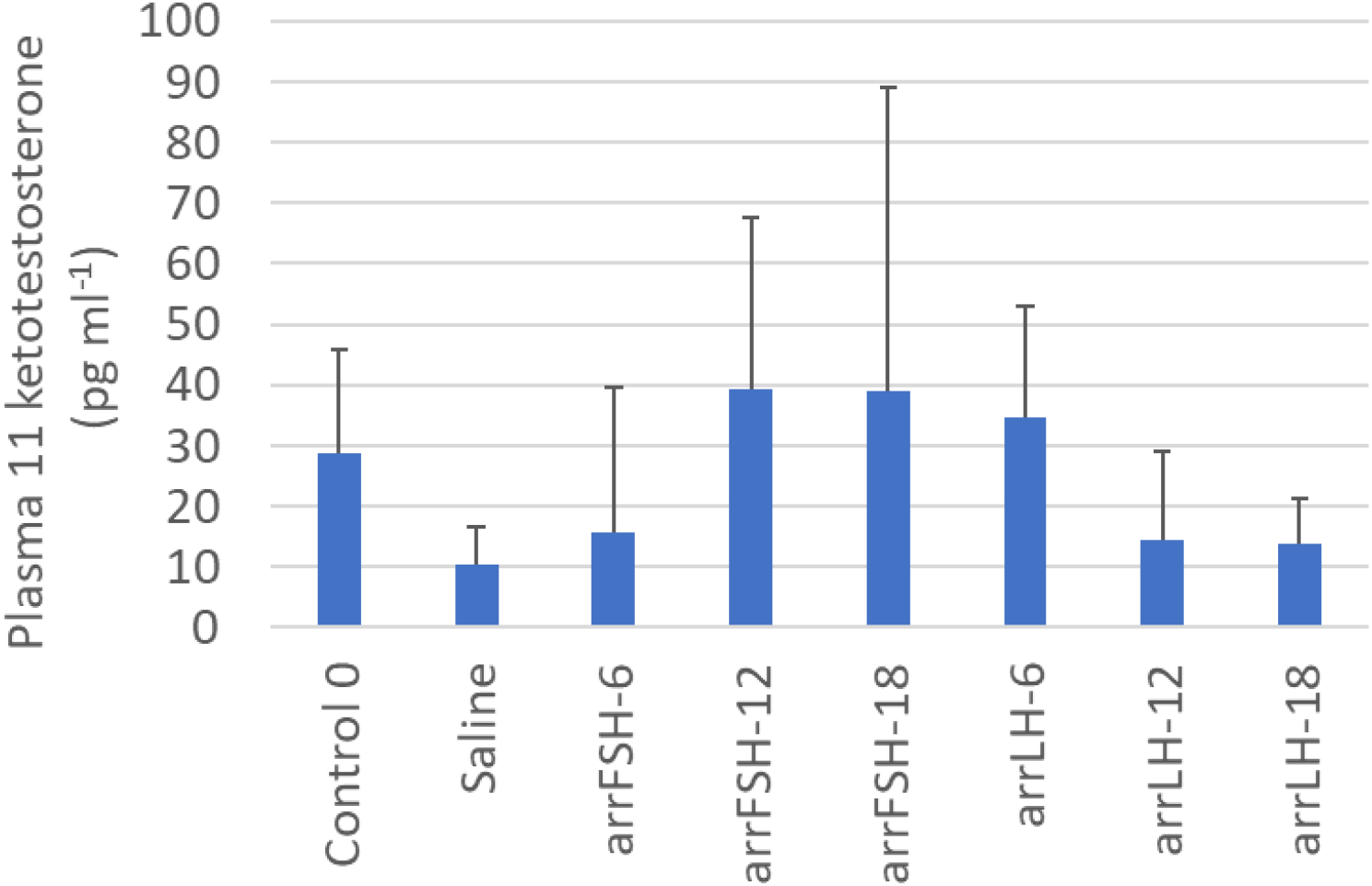
Mean plasma 11 ketotestosterone levels of meagre during sexual differentiation in arrGTH treatment groups and Control 0 at the start of the experiment (Week 0) and 4 days after the third weeks treatment (Week 3) of arrGTH or saline. Different letters indicate a significant (p<0.05) difference between week 0 and 3. The n for Control 0 was 61 and ranged from 6-8 for the different treatment groups.

**Figure 12.**
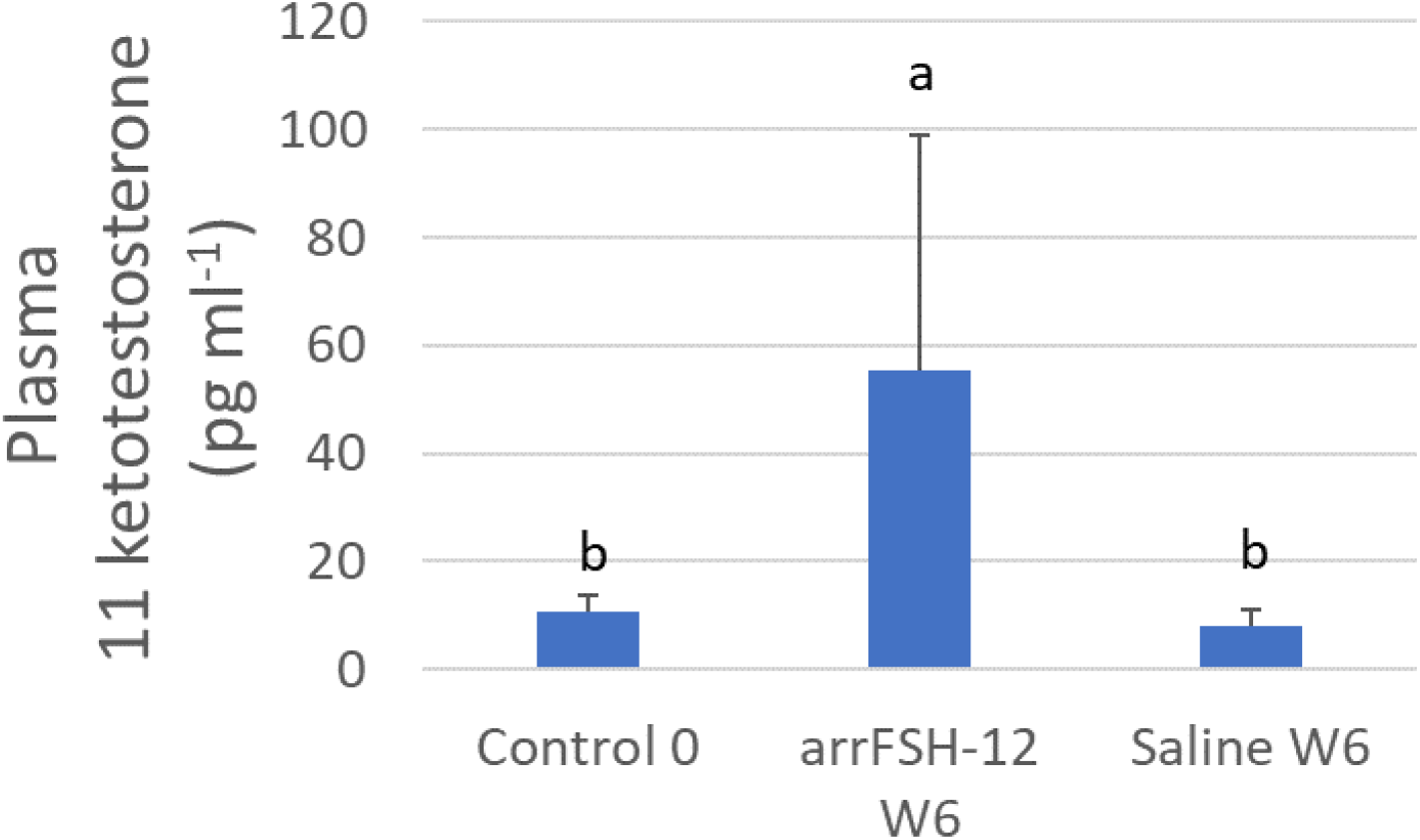
Mean plasma 11 ketotestosterone levels of meagre during sexual differentiation at the start of the experiment (Week 0) Control 0 and 7 days after the sixth weeks injection (Week 6) with 12 μg kg^-1^ of arrFSH (arrFSH-12 W6) or Saline (Saline W6 group). Different letters indicate a significant (p<0.05) difference between week 0 and 6. The n for Control 0 was 16 and 8 for groups arrFSH-12 W6 or SalineW6 group.

## 1.4 DISCUSSION

The arrGTH treatments did not induce gonadal development in meagre initiating sexual differentiation, although some development of isolated germ cells was observed. The gonads were classified as initiating sexual differentiation as histological cuts showed the space of the lumen or cavity in the centre of the gonad. The gonads contained either somatic cells or germ cells that were stained purple and orange. At the beginning of the experiment at time point zero the Control group had significantly higher E2 levels compared to three weeks later in both saline and arrGTH treated fish. The cell morphology and increase in E2 was similar to sexual differentiation described in meagre (Schiavone et al., 2012) and other species (Nagahama, 1999; Devlin and Nagahama, 2002). Sexually differentiating meagre had undifferentiated gonads at 5 months (October), increased E2 plasma levels at 12 months (March) and were differentiated with an ovarian lumen and oogonia or spermatogonia between 10 and 12 months (February to Mach) (Schiavone et al., 2012). This was similar to the more complete description in tilapia that initiated sexual differentiation with the production of E2 in females and the first morphological change was to develop an ovarian lumen and then the germ cells develop into oogonia, while males in the absence of E2 a testis lumen developed, and androgens increased as germ cells developed into spermatogonia (Nagahama, 1999; Kobayashi et al., 2013). These descriptions in meagre and tilapia indicate that the 9-month old (February) meagre in the present study were in the early stages of sexual differentiation exhibiting a gonadal lumen and increased E2 plasma levels, but almost no sexual differentiation of germ cells.

The arrGTH treatments had some minor effects on steroid production and GSI. The arrGTHs used in this study have been demonstrated to be biologically active. The same arrFSH preparations induced full spermatogenesis in 17 month-old, 1 kg meagre (Zupa et al., 2023) and applied to the same fish both arrFSH and arrLH were shown to increase steroid levels and induce spermiation (personal observation Neil Duncan). This is typical of the rGTHs produced with the CHO expression system that produce glycosylated proteins with high biological activity in vivo (Molés et al., 2020). In adult fish rGTHs have induced increases in steroid levels and spermatogenesis in Senegalese sole (*Solea senegalensis*) (Chauvigné et al., 2017, 2018), European eel (*Anguilla anguilla*) (Peñaranda et al., 2018), flathead grey mullet (*Mugil cephalus*) (Ramos-Júdez et al., 2021, 2022) and meagre (Zupa et al., 2023) and oogenesis in flathead grey mullet (Ramos-Júdez et al., 2021, 2022). However, despite of the demonstrated biological activity of the arrGTHs used in the present study, there was no effect of the arrGTHs on E2 plasma levels and actually plasma E2 decreased significantly in all treatments including control groups from week 0 to weeks 3 and 6. As mentioned this peak in E2 followed by a decrease was probably associated with the process of sexual differentiation (Nagahama, 1999; Schiavone et al., 2012; Kobayashi et al., 2013). There was no effect of arrGTHs on 11-KT plasma levels during the three-week experiment. The only significant difference observed in 11-KT was a significant increase in plasma 11-KT at 6 weeks in the treatment with 12 μg kg^-1^ of arrFSH. This increase appears to have been induced by the administration of arrFSH, which may have advanced the increase in 11-KT naturally associated with the sexual differentiation of males (Nagahama, 1999; Kobayashi et al., 2013). The GSI of fish at both 3 weeks and 6 weeks of treatment with 12 μg kg^-1^ of arrFSH had significantly lower GSI than fish at the start of the experiment (Control week 0), but were the same as saline Control fish GSI on weeks 3 and week 6. Similarly, to the steroids there was no effect of the arrGTHs on GSI.

In general terms there was also no effect of the arrGTHs on cell morphology. After three and six weeks of either arrFSH or arrLH at three different concentrations (over three weeks) the gonads had the same morphological appearance, with either somatic cells or undifferentiated germ cells. These cells accounted for over 99% of the cells in the gonads and the gonads after all treatments were classified as early stages of sexual differentiation with little change compared to week 0 and saline treated controls. However, the arrGTHs did have effects on the differentiation of isolated germ cells. The gonads presented a very low incidence, less than 0.1 % of differentiated germ cells. A significantly higher proportion of fish treated with arrGTHs had sexually differentiated germ cells. Male germ cells (type A and B spermatogonia, spermatocytes or spermatids) were the most common, but some fish had female perinucleolar oocytes and low proportion of fish were intersex with both male and female germ cells. A low proportion of Control fish also had isolated male germ cells, 2 fish from 16 (13%) compared to 25 % (2 fish from 8 treated with arrLH6) to 75% (6 fish from 8 treated with arrFSH6) of the fish treated with arrGTHs. This low abundance of isolated sexually differentiated germ cells may suggest that these germ cells and supporting somatic cells were at different stages of development compared to the majority of the germ cells in the gonads and these cells appeared to have had a direct or indirect capacity to respond to the application of arrGTHs and initiate an isolated development. It is not uncommon to observe isolated germ cell development at the initial stages of vitellogenesis (personal observation Neil Duncan) in the period when vitellogenesis is expected to initiate. As mentioned isolated male spermatogonia were observed in the Control fish, but the proportion of fish with isolated germ cell development was significantly increased by the arrGTHs indicating a minor, but significant effect.

Taken together the negligible effects of the arrGTHs, both arrFSH and arrLH on meagre initiating sexual differentiation have a range of interpretations that encompass (a) an aspect of the treatment was insufficient, for example the doses were not correct, (b) the gonadal stage was not receptive to the arrGTH treatments, for example the somatic cells were not at a developmental stage that enabled a response to the arrGTH signal, and (c) the GTHs do not have a role in meagre sexual differentiation. As previously mentioned the arrGTHs used in the present study have induced spermatogenesis (Zupa et al., 2023) and steroid production in 1 kg meagre (personal observation). In addition, rGTHs produced with the same expression system and at similar doses (12-18 μg kg^-1^) have successfully in adult fish induced spermatogenesis in Senegalese sole (*Solea senegalensis*) (Chauvigné et al., 2017, 2018), European eel (*Anguilla anguilla*) (Peñaranda et al., 2018), flathead grey mullet (*Mugil cephalus*) (Ramos-Júdez et al., 2021, 2022) and meagre (Zupa et al., 2023) and oogenesis in flathead grey mullet (Ramos-Júdez et al., 2021, 2022). These previous studies would indicate that the hormone and doses used were effective, although all these studies worked with fish at a more advanced stage of gametogenesis. Treatments with porcine FSH to orange-spotted grouper induced and accelerated sexual differentiation (Huang et al., 2019). A dose of 100 IU of porcine FSH accelerated differentiation into an ovary and then induced a sex change to testes. Interestingly in a short-term experiment, a low dose of 3 IU FSH upregulated gene expression of ovarian related genes and the highest does 100 IU upregulated gene expression of testes related genes. Huang et al., (2019) concluded that FSH was involved in sex differentiation in a concentration-dependent manner. This can be interpretated as an indication that the doses used in the present study may have been too high and had negative effects on the progression of sex differentiation. It is especially interesting that in the present study the isolated germ cell development was predominantly into male germ cells, which Huang et al., (2019) associated with high doses. The lowest FSH dose used in the present study 6 μg kg^-1^ of arrFSH (group arrFSH6) had the highest effect on progression of isolated germ cells into male germ cells, which may suggest that this low dose was a high dose for the sex differentiation process in meagre and that lower doses should be tested.

The possibility that the somatic cells were unable to receive and transduce the arrFSH signal should also be considered. At the stage of sexual differentiation, germ cells are associated with steroid-producing somatic cells that presumably receive an FSH signal in the gonads to produce the steroids, as Leydig and Granulosa cells were not developed during sex differentiation (Devlin and Nagahama, 2002). The steroid-producing cells would need to be present or at the correct stage of development to upregulate FSH receptors (FSHr) to receive and transduce the arrFSH signal. The European seabass has elevated FSH plasma during sexual differentiation (Moles et al., 2011) and FSHr and E2 changes during the first maturation (puberty) were similar with a weak correlation (Rocha et al., 2009) indicating the importance of a coordinated development with increased plasma FSH, upregulated FSHr and increased plasma E2. However, the higher plasma E2 at the start of the present study, would suggest that the steroid-producing cells were present to receive the arrFSH signal and suggests that as previously mentioned the signal was incorrect or the GTHs do not have a role in meagre sexual differentiation.

### 1.4.1 Conclusion

The role of the GTHs in sex differentiation has been questioned as immunoreactivity against FSHb and LHb (beta subunit of the gonadotropins) was not detected in the pituitary of the Malabar grouper during sexual differentiation (Murata et al., 2012) and the medaka phenotype with the FSH receptor knockout, completed sexual differentiation (Murozumi et al., 2014), both of which suggested that the GTHs did not play a major role in the sexual differentiation of these species. However, there is also growing evidence that GTHs are involved in sex differentiation as increases especially in FSH and related genes and receptors coincided with steroid production and sexual differentiation in different fish species (Fan et al., 2003; Moles et al., 2011; Fan et al., 2022) and the application of exogenous FSH induced sexual differentiation (Huang et al., 2019). In meagre, minor effects of the exogenous administration of arrFSH were observed, as arrFSH induced the development of isolated germ cells particularly male germ cells and a significant increase in 11-KT after 6 weeks of administration. These observations may indicate an involvement of FSH in the sexual differentiation of meagre as minor significant changes were observed in response to the exogenous application of arrFSH, but these minor changes need to be explained and this also highlight the gap in knowledge on the involvement the FSH in sexual determination and the need for more work to determine the period of sensitivity to exogenous FSH administration and the doses required.

## Acknowledgments

Thank you to the students Javier Gómez Aguilera and Lucas Stephen Arnold-Cruañes who helped generally with many aspects of the study. Thank you to Esteban Hernandez, Pol Moreno and Magda Monllaó for fish maintenance and sampling and Olga Bellot, Marta Sastre, Maria Corto and all IRTA technicians for lab work and sampling. Thank you to Ana Mendes, Pedro Pousão and IPMA, Olhão, Portugal for supplying the fish. Thank you to Marta Gut and Jèssica Gómez-Garrido from the Centre for Genomic Regulation (CRG), Barcelona, for sequencing of the GTH genes.

## Author contributions

Álvaro González Cid contribution was Data curation; Formal analysis; Investigation; Methodology; Validation; Visualization; Roles/Writing - original draft; and Writing - review & editing. Ignacio Giménez, contribution was Conceptualization; Funding acquisition; Investigation; Methodology; Project administration; Resources; Supervision; Validation; Visualization; Roles/Writing - original draft; and Writing - review & editing. Neil Duncan, contribution was Conceptualization; Data curation; Formal analysis; Funding acquisition; Investigation; Methodology; Project administration; Resources; Supervision; Validation; Visualization; Roles/Writing - original draft; and Writing - review & editing.

